# Retinal vascular dysfunction in the *Mthfr^677C>T^* mouse model of cerebrovascular disease

**DOI:** 10.1101/2024.12.31.630801

**Authors:** Alaina M. Reagan, Michael MacLean, Travis L. Cossette, Gareth R. Howell

## Abstract

**Introduction:** Investigating retinal biomarkers for Alzheimer’s disease (AD) and related dementias (RD), has increased significantly. We examine retinal vascular health in a mouse containing the ADRD risk variant *Mthfr^677C>T^*to determine if changes in retina mirror similar changes in cerebrovasculature.

**Methods:** Morphology and function of retinal vasculature and neurons were assessed using *in vivo* imaging, immunohistochemistry and pattern electroretinography. RNAscope and proteomics were employed to determine *Mthfr* gene expression and differential protein expression in mice carrying *Mthfr^677C>T^*.

**Results:** Mice show reduced retinal vascular density and reduced perfusion rate in aging mice, mirroring previously published brain data. *Mthfr* is widely expressed and colocalizes with vascular cell markers. Proteomics identified common molecular signatures across brain and retina.

**Discussion:** Results demonstrate that *Mthfr*-dependent vascular phenotypes occur in brain and retina similarly. These data suggest that assessing age and genetic-driven changes within retinal vasculature represents a minimally invasive method to predict AD-related cerebrovascular damage.

## 1. BACKGROUND

Determining reliable biomarkers for Alzheimer’s disease and related dementias (ADRDs) is critical for early diagnosis, increasing opportunities for intervention and treatment. While MRI and PET imaging provide critical information about brain function^1-4^ and cerebral spinal fluid can detect known markers of disease,^5,6^ the procedures can be invasive, painful, expensive, and inaccessible to many patients. Hence, these procedures are typically performed after a patient is symptomatic. Recent developments in blood biomarker discovery and detection are promising, and progress is being made regarding sensitivity and specificity.^7-9^ While blood biomarkers provide one component of clinical data, understanding early morphological and functional cellular changes in the brain is critical. Using the retina as a biomarker has been proposed and investigated,^10^ in part because of its accessibility in the clinic: retinal imaging occurs at most annual optometrist or ophthalmologist appointments. Additionally, because the retina is part of the central nervous system, it shares overlapping similarities to brain.^11,12^ Significant focus has been placed on identifying amyloid deposition in the retina,^13-15^ however, there is increasing evidence that additional retinal features, including changes within the vasculature has significant relevance to early ADRD detection.^16-19^

Cerebrovascular dysfunction is suggested to be one of the first perceptible changes before the onset of mild cognitive impairment.^20,21,22^ This dysfunction can occur through numerous mechanisms, including blood brain barrier disruption, hypoperfusion, hypertension, mini-bleeds, and small vessel disease.^23-25^ Because the brain and retina share similar cell types and the retinal vascular system experiences similar pressures to the cerebrovascular system,^26-28^ we predict that observable changes will occur in a parallel time frame to brain, potentially providing an opportunity for early detection. To test this prediction, this study utilized a mouse model carrying the common 677C>T variant in the methylenetetrahydrofolate reductase (*Mthfr*) gene to assess retinal vascular health as a predictor of cerebrovascular health. Genome-wide association studies have linked the 677C>T polymorphism (often referred to as *C677T*) (*MTHFR^677C>T^*) with Alzheimer’s disease^29-31^ and vascular dementia^32-34^ as well as with primary open angle glaucoma (POAG)^35,36^ and retinal vascular occlusive disease.^37^ ^38^ Up to forty percent of people carry at least one copy of the variant, so assessing its influence has broad relevance.^39,40^

We previously generated and characterized the *Mthfr^677C>T^*knock-in mouse model.^41^ The mouse phenocopies decreased liver MTHFR enzyme activity and increased plasma homocysteine. In addition, *Mthfr^677C>T^*mice showed cerebrovascular deficits by PET/CT, immunohistochemistry and electron microscopy that have been observed in Alzheimer’s disease and related dementias (ADRDs). These features make the *Mthfr^677C>T^* mouse the ideal model to test the biomarker potential of the retinal vasculature. We utilized *in vivo* ocular imaging and pattern electroretinography to assess baseline morphological features and function of the *Mthfr^677C>T^* retina as well as immunohistochemistry to confirm results. Proteomic analysis was performed to examine and compare differential expression in brain and retina. Data presented in the current study demonstrate that the presence of *Mthfr^677C>T^*results in reduced vascular density and increased risk for hypoperfusion, mirroring phenotypes in the cerebrovasculature and highlighting its use as a predictive tool for brain pathology. Differential protein expression, shared between brain and retina from *Mthfr^677C>T^* mice, suggests the variant alters pathways involving metabolism and cell survival similarly in both tissues.

## 2. METHODS

### 2.1. Ethics statement

All research was approved by The Jackson Laboratory Institutional Animal Care and Use Committee (IACUC; approval number 12005). Animals were humanely euthanized with 4% tribromoethanol (800 mg/kg). Authors performed their work following guidelines established by the eighth edition of the *Guide for the Care and Use of Laboratory Animals* and euthanized using methods approved by the American Veterinary Medical Association.

### 2.2. Mouse Strains

The *Mthfr^677C>T^* mouse strain (JAX ID: 031464) was generated at The Jackson Laboratory (JAX) as part of the IU/JAX/Pitt/Sage MODEL-AD Center under the MODEL-AD animal use summary (ID:18051) and maintained under the Howell lab animal use summary (ID: 12005). Experiments were conducted in accordance with policies and procedures described in the Guide for the Care and Use of Laboratory Animals of the National Institutes of Health^42^ and were approved by the JAX Institutional Animal Care and Use Committee. All mice are congenic on the C57BL/6J (JAX# 000664) (B6) strain and bred and housed in a 12/12-hour light/dark cycle and fed LabDiet 5K52 (LabDiet), a standard natural ingredient diet containing 1.9 mg/kg folic acid, and 9.0 mg/kg riboflavin. In mice, based on numbering from ENSMUST00000069604.15, the 806C>T causes a codon change GCC>GCT, leading to an A262V mutation in the methylenetetrahydrofolate reductase gene that is equivalent to the 677C>T polymorphism and corresponding A222V mutation in humans. Given its common usage, we refer to this mouse model as *Mthfr^677C>T^*. To produce experimental animals, mice heterozygous for *Mthfr^677C>T^*(*Mthfr^C/T^)* mice were intercrossed to produce litter-matched male and female *Mthfr^C/C^*, *Mthfr^C/T^* and *Mthfr^T/T^* mice (herein referred to as CC, CT and TT). Mice were aged to either 6 months or 12 months.

### 2.3. Optical coherence tomography, fundus, and fluorescein angiography

Optical coherence tomography (OCT) fundus exams, and fluorescein angiographies were performed using Micron IV and Reveal OCT and Discover 2.4 software (Phoenix-Micron, Bend, OR). Mice were given 1 drop (20-30ul) of 0.5% Tropicamide (Somerset Therapeutics, NDC# 70069-121-01) in both eyes. After 10min, one drop (20-30ul) of 2.5% phenylephrine (Bausch & Lomb, NDC# 42702-0103-05) was applied to both eyes. Mice were then anesthetized using 4% isoflurane until a proper plane of anesthesia is achieved. Mice were then transferred to a nose cone on a heated holding cradle. Anesthesia was then reduced to 2% isoflurane. Eyes were kept moist using GenTeal Tears Lubricant Eye Gel Drops (Alcon). Respiration rate was monitored, and the camera lens was adjusted to be perpendicular to the cornea.

For OCT imaging, focus and brightness of the image guided OCT was adjusted until optimal image is previewed. OCTs were captured from the temporal to the nasal retina in a plane that included the optic nerve. To improve resolution, 10-40 images were taken and averaged. For fundus exams, a white light fundus image was taken of each eye. Brightness adjustments were made, and the focal plane was optimally adjusted. Immediately following the image guided OCT and fundus imaging, mice were injected intraperitoneally (IP) with 1% Fluorescite® Fluorescein Sodium at a dose of 10mg/kg (10mg/mL solution, Akorn). Using a GFP filter, images of the right eye were taken every 30 seconds for 6 minutes. Images were taken of the left eye 30 seconds after the right eye imaging period.

Retinal layers from OCT images were measured using the FIJI distribution of ImageJ (46,47). Each layer was measured 200, 400, and 600µm from the edge of the optic nerve using the line measurement tool, and the 3 respective measurements were averaged and used for statistical analyses. (n=6/sex/genotype)

### 2.4 Mouse Perfusion and Tissue Preparation

Mice were anesthetized with a lethal dose of Tribomoethanol based on animal mass and transcardially perfused with 1X PBS (Phosphate Buffered Saline). Liver blanching was used to determine a successful perfusion. Retina and brain tissue to be used for proteomic assessment were quickly removed and snap frozen in liquid nitrogen. For immunohistochemical assessment, brain hemispheres and eyes were fixed in 4% Paraformaldehyde (PFA) overnight at 4°C. Retinas were then dissected and prepared for the respective staining protocols.

### 2.5. Immunostaining

Retinas were placed in 1X PBT (PBS + 1% Triton 100X) for 1 hour at room temperature, followed by blocking in 10% normal donkey serum overnight at 4°C. Retinas were then incubated in primary antibodies diluted with 1X PBT + 10% normal donkey serum overnight at 4°C. After incubation with primary antibodies, sections were rinsed three times with 1X PBT for 15 min and incubated overnight at 4°C in the corresponding secondary antibodies. Tissue was then washed three times with 1X PBT for 15 min, incubated with DAPI for 5 minutes, washed with 1X PBS and mounted in Poly aquamount (Polysciences). The following primary antibodies were used: goat anti-Collagen IV (ColIV) (1:40, EMD Millipore Cat# AB769) and rat anti-CD31 (1:100). Secondary antibodies: donkey anti-goat 488 (1:500, Invitrogen), donkey anti-goat 647 (1:500 Invitrogen) donkey anti-rat 594 (1:500 Invitrogen). Antibodies listed in this study have previously been validated in the Howell lab and are frequently used and maintained (n=6/sex/genotype).

### 2.6 Pattern electroretinography

All pattern electroretinography (PERG) was performed using a JORVEC PERG system (Intelligent Hearing Systems, Miami, Florida) as described by the manufacturer. Briefly, mice were anesthetized with ketamine/xylazine, and slit lamp was used to inspect each mouse eye prior to starting experiment. Mice were kept on a warming stage throughout the experiment. Electrodes were placed subcutaneously such that the active electrode is placed between the eyes with the tip just before the snout and the reference electrode is placed in line with the active electrode just between the ears. The ground electrode was placed approximately 2cm in front of the base of the tail. Scans were collected and waveforms folded using default settings. PERG amplitude was calculated as P1-N2 for each eye per mouse (n=6/sex/age/genotype).

### 2.7. RNAscope

Brain hemispheres and eyes were collected following PBS perfusion and fixed in 10% Neutral Buffered Formalin overnight before being transferred to 70% ethanol for 24 hours and embedded in paraffin after gradual ethanol dehydration. Coronal sections were cut at 5 μm. For brain, every tenth section was mounted onto 2 slides, with three sections on each slide. Sections were stored at room temperature. RNAscope was performed using the ACD RNAscope LS 4-plex Fluorescent Assay and RNAscope LS 2.5 Probes for Mm-Tagln (Cat# 480338-C2), Mm-Pecam1 (Cat# 316728-C3), Mm-Pdgfrb (Cat#411388-C4), and Mm-Mthfr (Cat#839028). The slides were placed on the Leica Bond for automated deparaffinization and blocking in hydrogen peroxide for 10 min at room temperature and pretreated using Target Retrieval Solution at 95 °C for 20 min and Protease Plus at 40 °C for 30 min. Probe hybridization and signal amplification were performed according to the manufacturer’s instructions for fluorescent assays. TSA vivid fluorophores 520, 570 and 650 were used in combination for probe visualization at a 1:1500 dilution. The slides were imaged on an SP8 confocal (Leica). Transcripts were assessed under 20x and 40x magnification in representative regions (n=3/sex/genotype).

### 2.8 Proteomic Analysis

#### Protein Extraction and Sample Preparation

Snap-frozen plasma (20 uL), retina (one per mouse), and brain samples from a cohort of 36 *Mthfr^677C>T^* mice (n=6/sex/genotype) were provided to the Mass Spectrometry and Protein Chemistry Service at The Jackson Laboratory for dual proteomics and metabolomics extraction.^43,44^ 600 µL of ice-cold extraction buffer (2:2:1 methanol:acetonitrile:water) plus internal standards (0.5 ng/uL caffeine, 1-NAP, 9ANC) were added to each 20 µL plasma and retina sample. Brain samples all had an equal ratio of 2:2:1 extraction buffer added with 200 mg set to 1 mL of buffer and scaled appropriately. Retina and brain samples also had a pre-chilled 5 mm stainless steel bead (QIAGEN) added to the tube. Sample tubes and Tissue Lyser II cassettes were pre-chilled at -20°C. Batches of 48 samples were then lysed in the Tissue Lyser II for 2 minutes at 30 1/s; assessed and repeated for 1 minute only if necessary. Stainless steel beads were removed with a magnet and the samples were extracted overnight for 16 hours at -20°C. Sample extracts were centrifuged at 21,000 x g at 4°C for 15 minutes. The supernatant (containing metabolites) was removed and put in the -80°C for future metabolomics analysis. The protein pellet was reconstituted in 50 mM HEPES buffer, pH 8.2, containing 6M Urea (100 uL for retina samples, 500 uL for brain samples, 250 uL for plasma samples). Reconstituted samples were vortexed at max speed for 30 seconds, a chilled steel bead was added, and they were reconstituted using the Tissue Lyser II as performed in the extraction step. Samples were then ice-waterbath sonicated (sweep at 37 Hz at 100% power) for 5 minutes (30 seconds on, 30 seconds off for five cycles). Samples were assessed for clarity and the sonication was repeated only if necessary. Samples were centrifuged at 21,000 x g for 15 minutes at 4°C to pellet any heavy debris and protein supernatant was transferred to a new 1.5 mL microcentrifuge tube. Samples were then snap-frozen and stored at -80°C until further use.

#### Protein Digestion and Peptide Purification

Protein lysates were quantified using a microBCA assay (Thermo, Cat.# 23235) according to manufacturer protocol. For all samples from each tissue 20 µg of each sample were aliquoted and brought to an equal volume in 50 mM HEPES, pH 8.2 to dilute the Urea to <2M final concentration. Samples were reduced with 10 mM dithiothreitol at 37°C for 30 minutes with 500 rpm agitation (ThermoMixer), alkylated with 15 mM iodoacetamide for 20 minutes at room temperature with 500 rpm agitation (ThermoMixer), and trypsin-digested (Sequence Grade Modified; Promega) with a 1:50 trypsin:protein ratio at 37°C for 20 hours with 500 rpm agitation (ThermoMixer). Digest peptides were then clean-up using Millipore C18 zip-tips (Millipore, Cat. #ZTC18S096) according to the manufacturer protocols.^45,46^ Eluted peptides were dried using a vacuum centrifuge and stored at -20°C until reconstitution for tandem mass spectrometry analysis.

#### Liquid Chromatography Tandem Mass Spectrometry Analysis (LC-MS/MS)

Each of the dried peptide samples were reconstituted in 20 µL of 98% H2O/2% ACN with 0.1% formic acid via pipetting and vortexing. Reconstituted samples were then transferred to a mass spec vial and placed in the autosampler at 4°C. Tandem mass spectrometry analysis was then performed on a Thermo Eclipse Tribrid Orbitrap with a FAIMS coupled to an UltiMate 3000 nano-flow liquid chromatography system using an EASY-Spray column (PepMap RSLC C18, 2 µm, 100 Å, 75 µm x 50 cm) in the Mass Spectrometry and Protein Chemistry Service at The Jackson Laboratory; each sample injection consisted of 8 uL of reconstituted sample and a technical replicate run was run for each. The method duration was 90 minutes using Buffer A (100% H2O with 0.1% formic acid) and Buffer B (100% acetonitrile with 0.1% formic acid) with a flow rate of 300 nL/min. The full gradient consisted of 98% A/2% B from 0-5 minutes, 98% A/2% B at 5 minutes to 92.5% A/7.5% B at 10 minutes, 92.5% A/7.5% B at 10 minutes to 70% A/30% B at 70 minutes, and 70% A/30% B at 70 minutes to 10% A/90% B at 75 minutes. The gradient was held at 10% A/90% B until 78 minutes and brought to 98% A/2% B by 80 minutes, where it was held for 10 minutes to equilibrate the column. Method settings included a default charge = 2, expected peak width = 30 seconds, advanced peak determination, spray voltage = 2000V, mode = positive, FAIMS carrier gas =3.5 L/min, an ion transfer tube temperature of 325°C, and FAIMS voltages of -40V and -50V. Settings for MS1 precursor spectra detection included a cycle time = 1 second (each node), Orbitrap resolution of 120,000, scan range from 375-1200 m/z, RF lens at 30%, normalize AGC target of 250%, maximum inject time (ms) = auto, microscans = 1, data type = profile, monoisotopic precursor selection = peptide, minimum intensity threshold of 5.0e3, charge states = 2-7, and a dynamic exclusion of a n = 1 for 60 seconds. MS2 peptide fragment analysis was performed in the ion trap with an isolation window of 0.7 m/z, a fixed collision energy of 33% using HCD activation, ion trap scan rate = turbo, maximum inject time = 10 ms, normalized AGC target of 100%, and centroid data settings.

#### Tandem Mass Spectrometry Data Analysis

The Thermo Eclipse RAW data files from the Eclipse Tribrid Orbitrap for each sample were searched against the UniProtKB Mus musculus (sp_tr_incl_isoforms TaxID=10090; v2021-02-04) protein database in Proteome Discoverer (version 2.5.0.400) using Sequest HT according to standard manufacturer recommended workflows. The Sequest protein database search parameters included trypsin cleavage, precursor mass tolerance = 20 ppm, fragment mass tolerance = 0.5 Da, dynamic carbamidomethylation of cysteine (+57.021 Da), dynamic methionine oxidation (+15.995 Da), and dynamic N-terminal acetylation modification (+42.011 Da). Other setting included a maximum modifications per peptide of 3, a maximum number of missed cleavages of 2, a minimum peptide length of 6 amino acids, a maximum peptide of 144 amino acids, and a fragment mass tolerance = 0.6 Da. Additional thresholding settings included the use of the Percolator node, concatenated target/decoy selection, q-value validation, a maximum delta Cn of 0.05, and a false discovery rate of less than 0.05 for all matches. The Thermo recommended default Minora node parameters were also utilized. Abundance values were normalized to the total ion signal in the samples and pairwise ratio-based peptide analysis was used for the summed abundances. Strict parsimony principle was then applied for protein grouping and master proteins were reported.

#### Differential Proteomics Analyses

All analyses were completed within each tissue (retina, brain, and plasma) separately. Identified peptides with FDR<0.05 confidence and detected in at least three samples were used for further investigation. log2(scaled abundance+1) values were used for differential expression analyses. Differentially expressed proteins (DEPs) were determined using generalized linear models and a repeated measures model via limma and DEqMS^47,48^ which takes into account the number of unique peptide spectrum matches (PSMs) used for each peptide to enhance the statistical power of differential expression, with either 1) groups defined as a combination of genotype and sex (∼0+Group) and contrasts designed to test for genotype effects while controlling for any sex differences. As samples were run in duplicate, we utilized the duplicateCorrelation function within limma to block on “mouseID”.^48^ clusterProfiler v4.6.2 R package was used to test for overrepresentation of Gene Ontology (GO) Biological Process gene sets within the list of differentially expressed proteins common to both the brain and retina between genotypes with a FDR of less than 0.05 and enrichPlot v1.18.4 were used to visualize the enrichment results.^49-55^ STRING-dB v 2.14.3 R package was used to visualize protein-protein interaction networks with the settings “score_threshold=200” and “network_type=physical” and to calculate enrichment of GO gene sets within the list of DEPs.^50,56^

### 2.9. Imaging analysis

Fiji (version 2.14.0/1.54f) was used for quantification of vessel area in *in vivo* fluorescence angiography images. The “Vessel Analysis” plugin was used to automatically calculate vascular density metrics: vascular density = vessel area/total area * 100% - vascular length density = skeletonized vessel area/total area * 100%. in both *in vivo* and *ex vivo* images, the circle tool was used to determine a fixed distance from the optic nerve head in which to measure vessel diameter. The measuring tool was then used to calculate individual arteriole and venule diameters. All measurements are represented graphically.

Z-stacks of immunostained retinal wholemounts were taken on a confocal microscope (Leica TCS Sp8 AOBS with 9 fixed laser lines) at 20x magnification using the Leica Application Suite X (LAS X) software (Leica Microsystems GmbH, Mannheim, Germany). Images were taken at the optic nerve head and the retinal periphery (*n*=6/sex/genotype/age for a total of 216 images).

### 2.10. Statistical Analysis

Sample size was determined to be sufficient using previously published work from the Howell lab as well as previously published work from our collaborators**.^57-60^** Because retinal tissue was required for each assay, separate cohorts of mice had to be developed, maintained and aged to perform experiments. For all experiments, authors were blinded to animal genotype before data generation. Data are shown as the mean ± standard deviation. Normality was assessed using Shapiro-Wilk and Kolmogorov-Smirnov tests. Multiple group comparisons were performed by one-way or two-way multifactorial analysis of variance (ANOVA) depending on the number of variables followed by Tukey post hoc test. If data from multiple group comparisons was not normally distributed, the Kruskal-Wallace tests were used followed by the Dunn’s multiple comparisons post hoc test. When assessing differences between two groups, a paired two-tailed Student’s test was used unless the data were not normally distributed and a Mann-Whitney test was performed. Differences were considered to be significant at *p* < 0.05. Statistical analyses were performed utilizing Prism v9.3.1 software (GraphPad Software Inc., San Diego, CA, USA). For all analyses, * p<0.05, ** p<0.01, *** p<0.001, **** p<0.0001. Analysis for proteomic data is included in section 2.4.

## 3. RESULTS

### 3.1. *Mthfr^677C>T^* retinas show age- and sex-dependent reduction of vascular density

*In vivo* fluorescein images were used to examine prominent retinal vasculature throughout the anterior layer of the tissue. A Fiji plugin was used to analyze images and measure vascular density (Figure 1A). We have previously shown a reduction in cortical vessel density in *Mthfr^677C>T^* mice, with female mice showing a more pronounced reduction.^41^ Similarly, we observe a reduction in total retinal vascular density in 12-month female TT mice when compared to male mice of either age and 6-month female TT mice, suggesting that the TT genotype has more effect with age and in females. (Figure 1B) Additionally, we show reduced numbers of total branching vessels in 12-month female TT mice, specifically suggesting a simplification of the large-vessel network in the retina (Figure 1C) (n=6/sex/age/genotype).

**Figure 1.**
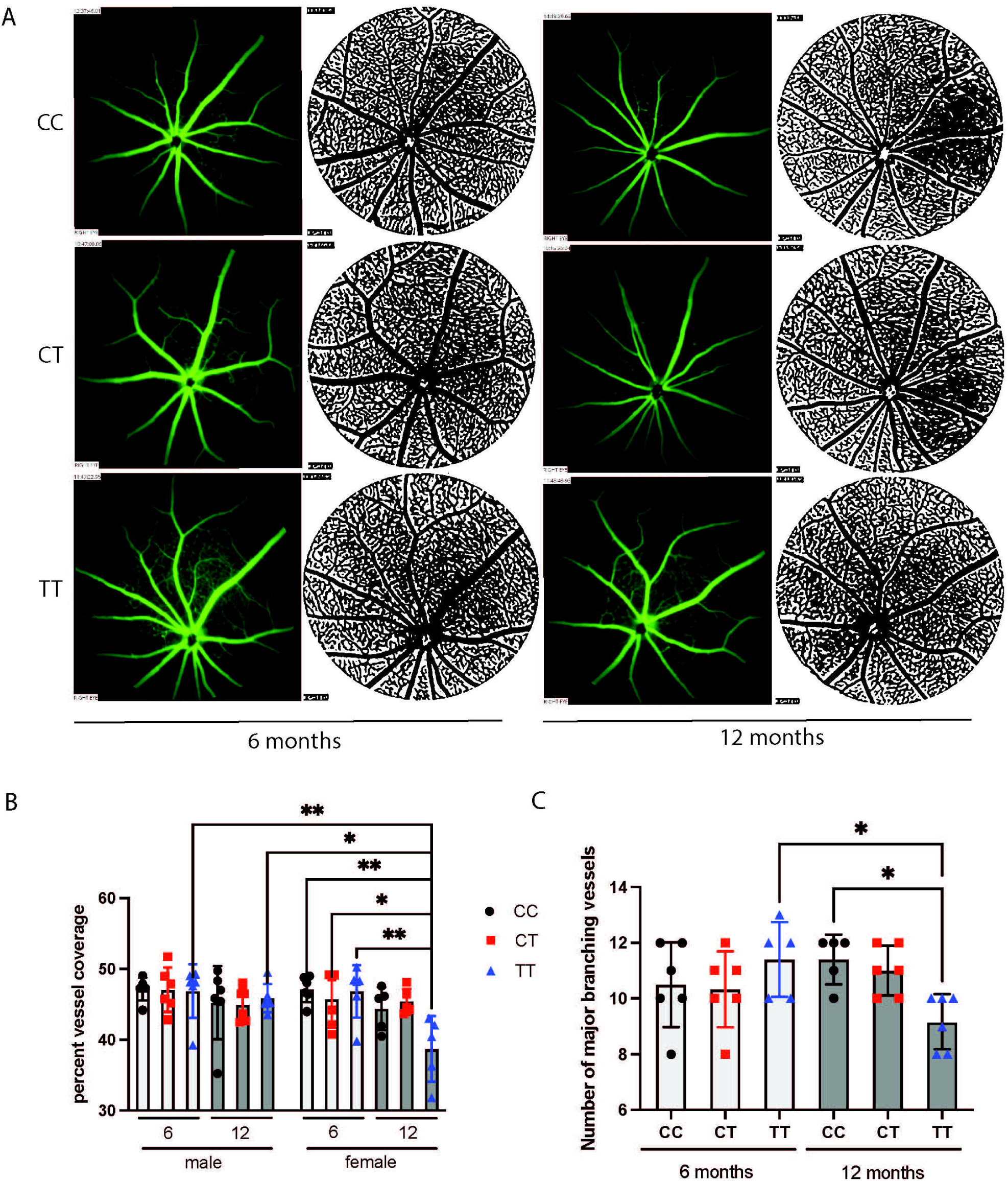
*Mthfr^677C>T^* retinas show age- and sex-dependent reduction of vascular density. Fluorescence angiography was used to take *in vivo* images of the retinal vasculature. Post-fluorescein injection, retinas were imaged every minute for six minutes, and images used for analysis were taken at the 4-minute mark from each animal. A Fiji plugin was used to measure vascular density. (Figure 1A) Quantification of the vascular density shows reduced vascular density in aged female TT mice. (Figure 1B) The number of major branching vessels was also reduced with age in male and female TT mice. No significant differences in branching number were detected between males and females within age/genotype groups, so numbers were averaged for quantification (Figure 1C). n=6/sex/age/genotype.

### 3.2. *In vivo* and *ex vivo* imaging suggests altered blood perfusion in 677C>T mice

At baseline, retinal fundus images from mice carrying the 677C>T variant do not show abnormalities at 6 months of age (Figure 2A left panel). Following an IP fluorescein injection, images of the right eye were taken every 30 seconds for 6 minutes using a GFP filter. Representative images were taken at the 4-minute mark (Figure 2A right panel). Using the value of the fluorescein dye intensity over the imaging time course, we were able to approximate the rate of the blood flow into the retina. We detected differences in the mean intensity of dye across genotypes and ages in males (Figure 2B) and females (Figure 2C). Interestingly, age reduced the mean intensity across all CC and TT mice. CC males and females had a 14% and 11% reduction in mean fluorescence intensity of fluorescein with age, respectively. TT males and females had a 9% and 13% reduction with age, respectively. TT genotype also showed a reduced intensity at baseline compared to CC, with a reduction of 11% and 6% in 6-month males and females, respectively. 12-month TT mice have a 5% (males) and 6% (females) reduction in intensity compared to 12-month CC mice, with a reduction of 19% and 18% intensity overall when comparing 6-month CC to 12-month TT, suggesting a subtle, but relevant genotype-dependent reduction in blood perfusion rate in the retina.

**Figure 2.**
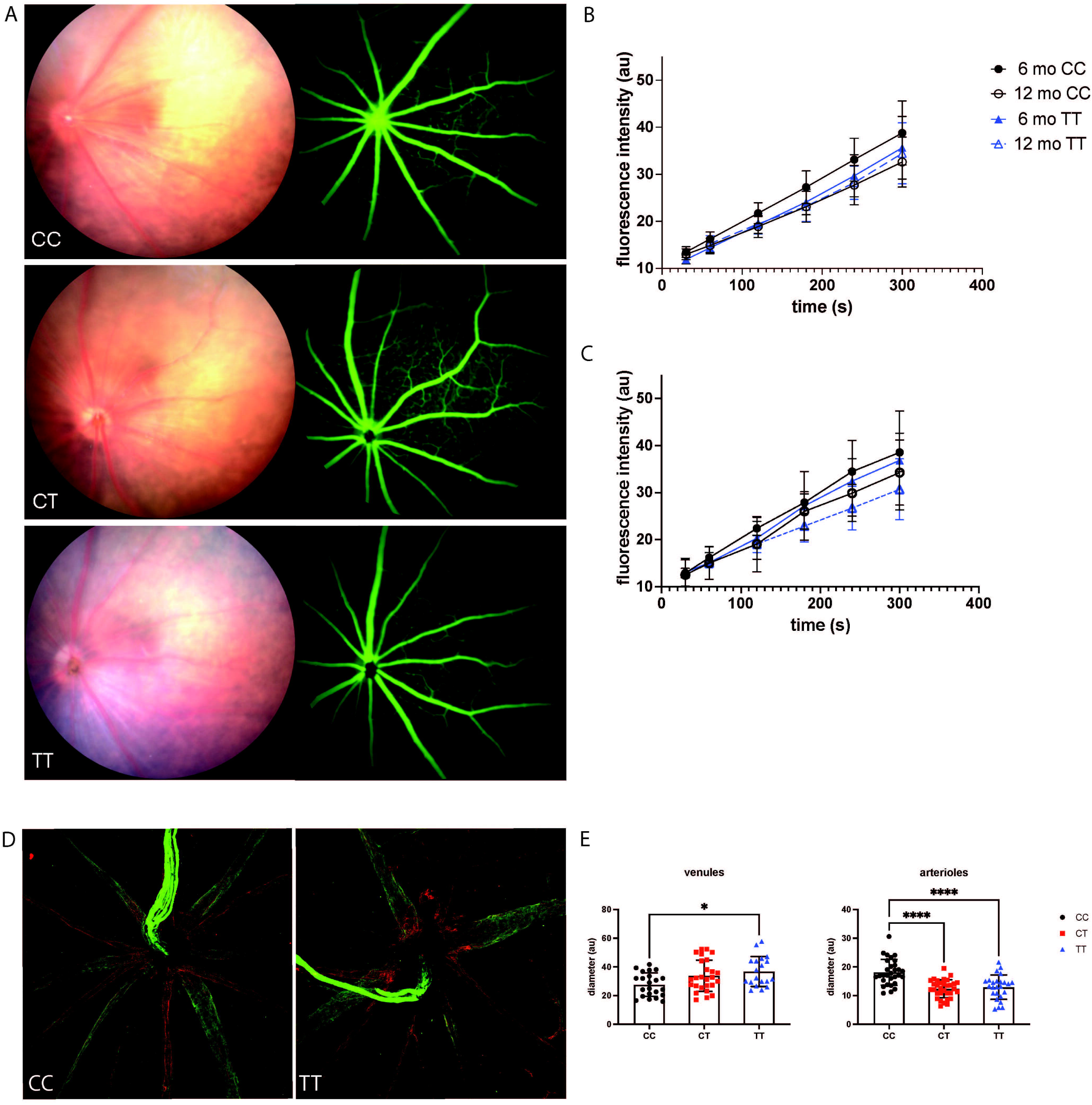
*In vivo* and *ex vivo* imaging suggests altered blood perfusion in 677C>T mice. Representative fundus (Figure 2A left panel) and fluorescent angiography (Figure 3A right panel) images from 6-month mice of each genotype. The rate of fluorescein dye intensity was measured over five minutes to approximate of rate of blood flow. CC male mice showed a decrease in fluorescent intensity with age, while TT male mice showed reduced intensity by 6 months compared to age-matched controls (Figure 2B). Both CC and TT female mice showed decreased fluorescent intensity with age, with TT mice showing a genotype-specific reduction overall (Figure 2C) n=6/sex/genotype. Ex vivo, retinal wholemounts were immunostained for collagen IV (green) and CD31 (red). All branching arterioles and venule diameters were measured from a set distance from the optic nerve head. Representative images (Figure 2D) show reduced overall arteriole diameter and increased venule diameter (Figure 2E) n=6/sex/genotype; number of total vessels differed between mice.

While overall vessel density was reduced in 12-month TT females, (Figure 1B) this analysis does not account for changes in individual vessel types. Diameters of arterioles and venules near the optic nerve head were assessed individually in 6-month mice; the presence of *Mthfr^677C>T^* results in arterial narrowing (Figure 2D) and venous enlargement (Figure 2E). (n=6/sex/genotype).

### 3.3. Hypertensive-like phenotypes are observed in *Mthfr^677C>T^* retinas

Clinically, retinal arteriovenous crossing (AVC) is observed in patients with ocular disease^61^ and/or vascular disease^60,62^ and consists of an arteriole lying across a venule, decreasing blood egress on the venous side. we examined the fluorescein images for the AVC phenotype. AVC occurred in all animals, but the percent of events increased in female CT and TT mice at 6 and 12-months and in TT males at 12-months (Figure 3A-B).

**Figure 3.**
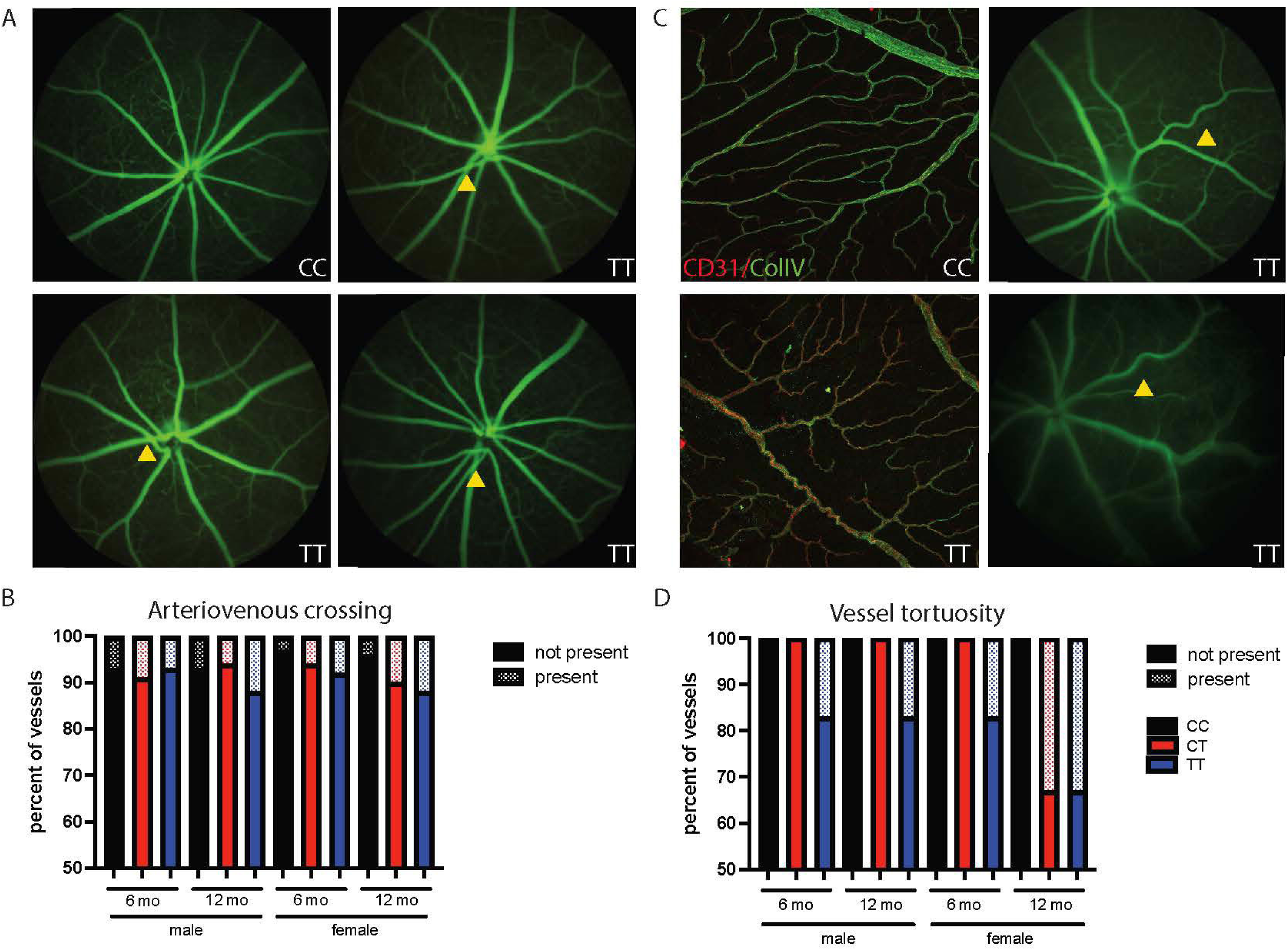
Human disease-relevant vascular phenotypes are observed in *Mthfr^677C>T^* retinas. Representative fluorescein angiography images show healthy retinal vasculature with even spacing between arterioles and venules in a CC mouse (Figure 3A top left). Representative images showing examples of arteriovenous crossing in TT mice (Figure 3A top right, bottom left and right; yellow arrowheads). The number of arteriovenous occurrences were counted in each retina and averaged per group for a total percent of vessels with occurrences. Mice carrying the variant showed an increase in occurrences, with aged TT mice showing the highest percents (Figure 3B). Representative retinal wholemounts immunostained for antibodies against CD31 (endothelial) and collagen IV (extracellular matrix) show normal vessel patterning in CC mice (Figure 3C, top left panel), TT mice show increased vessel tortuosity (Figure 3C, bottom left panel.) In vivo, tortuous vessels can be observed in TT mice (Figure 3C, right panels). When in vivo images were quantified, occurrences of tortuous vessels occurred only in TT mice at 6 months of age and increased in number with age and in female mice (Figure 3D).

Retinal vessel tortuosity also occurs in patients with ocular disease and/or vascular disease and presents as twisted vessels. This can result in turbulent blood flow, which promotes changes in blood pressure, degradation of elastin and luminal shear stress (Han 2012). Tortuous vessels were observed in both *ex vivo* immunostaining (Figure 3C, left panels) and *in vivo* angiography (Figure 3C, right panels). Specifically, we observed an increased number of tortuous events in all TT mice compared to CC, but specifically in both CT and TT 12-month females (Figure 3C-D) (n=6/sex/genotype).

### 3.4. *Mthfr^677C>T^* retinas do not demonstrate neuronal morphological or functional deficits at baseline

To investigate overall morphological changes in the *Mthfr^677C>T^*retinas, OCT was performed to examine genotype-driven changes in retinal thickness. An increase in retinal thickness can represent inflammation or edema, while retinal thinning represents neuronal loss. Retinal thickness was measured at three set distances from the optic nerve head (Figure 4B) and averaged for analysis (Figure 4C). The overall thickness of retinas from all groups do not show any genotype, sex, or age-related changes at baseline.

**Figure 4.**
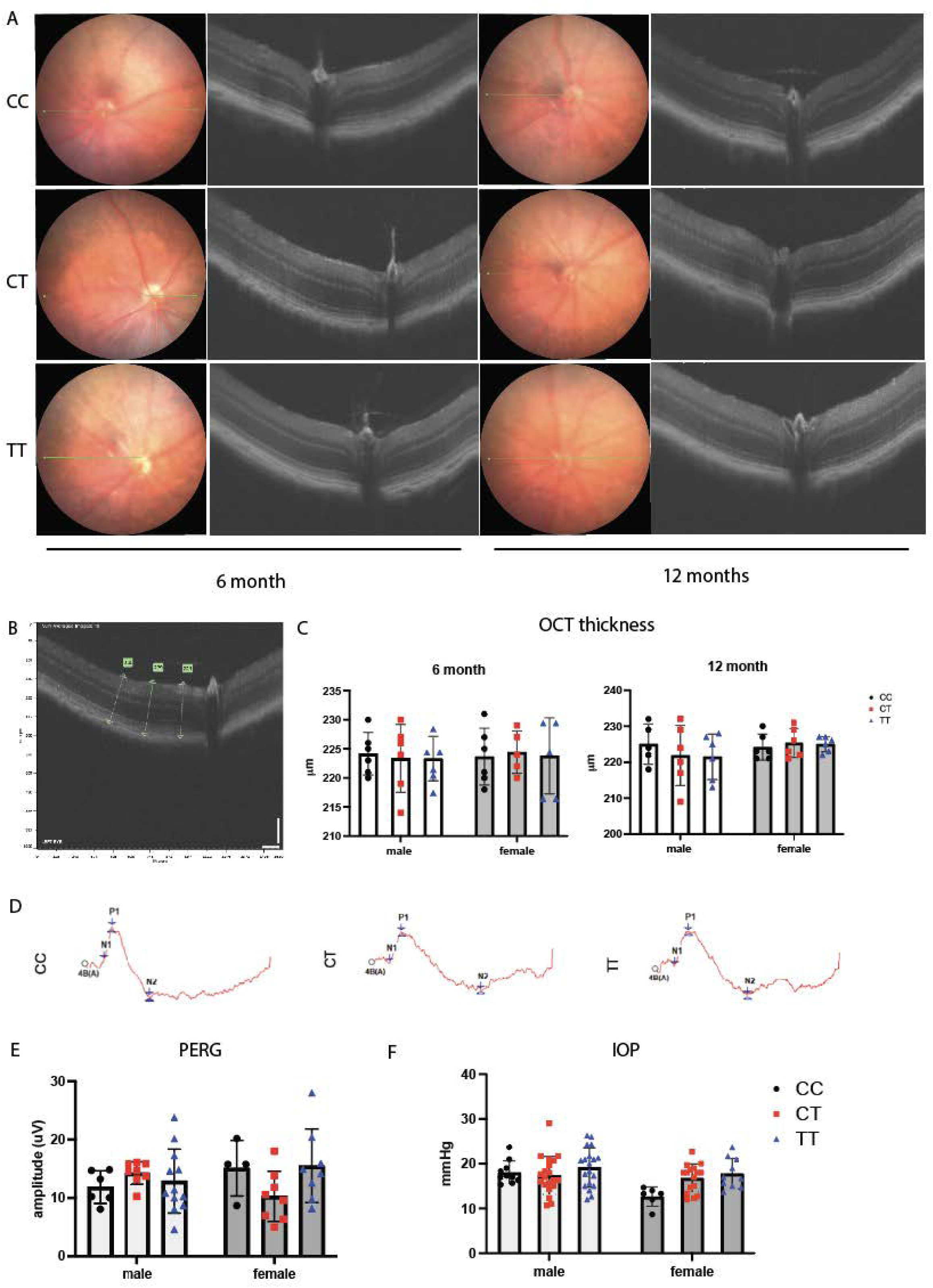
*Mthfr^677C>T^* retinas do not demonstrate neuronal morphological or functional deficits at baseline. Representative fundus and OCT images from mice of all genotypes at 6 and 12 months of age (Figure 4A). Retinal thickness was measured at 3 distances from the optic nerve head (Figure 4B) and averaged for quantification. Mice do not show genotype-specific differences in thickness (Figure 4C). n=6/sex/genotype/age. Representative traces from PERG measurement (Figure 4D) highlight that no significant genotype-specific differences are seen in waveform amplitude at 6 months of age (Figure 4E). Mice also do not show differences in IOP at 6 months (Figure 4F). n=6/sex/genotype

Retinal ganglion cells (RGCs) are the primary neurons damaged in glaucoma. Because there is evidence that *Mthfr^677C>T^* is a risk factor for glaucoma, we assessed RGC function using pattern electroretinography (PERG) to probe for gross changes in RGC potential. Representative PERG traces (Figure 4D) and analysis (Figure 4E) show no significant differences in signal amplitude in 6-month mice. Elevated intraocular pressure (IOP) is a major risk factor for glaucoma, so IOP was measured in *Mthfr^677C>T^*mice, but also showed no significant differences between sex or genotype (Figure 4F). Collectively, these data suggest the variant alone is not inducing retinal neurodegeneration (Figure 4A) (n=6/sex/genotype).

### 3.5. *Mthfr* is widely expressed in the retina and brain

Data presented here and our previously published work (me) show overlapping phenotypes between the retinal and cerebral vasculature in *Mthfr^677C>T^* mice. To begin to determine if the mechanisms underlying these changes are similar, we first assessed *Mthfr* expression in retina and brain. Existing bulk (Barres) and single cell (Vanlandewijck and He) RNA sequencing datasets show *Mthfr* expression in numerous cells in mouse and human brain, including astrocytes, neurons, oligodendrocytes, microglia/macrophages, endothelial cells, pericytes, and smooth muscle cells. To compare *Mthfr* expression in retinal tissue with brain tissue *in situ*, we performed RNAscope on 6-month CC and TT mice. We observed widespread expression of *Mthfr* in retina (Figure 5A) and brain tissue (Figure 5B-C), with *Mthfr* expression colocalizing with probes for *Pecam1*, *Pdgfrβ* and *Tagln*, which are known to be expressed in endothelial cells, pericytes and smooth muscle cells, respectively. Here expression appears to be distributed in a similar manner in both tissues regardless of genotype, supporting the use of the model to study parallel vascular events. (n=3/sex/genotype.)

**Figure 5.**
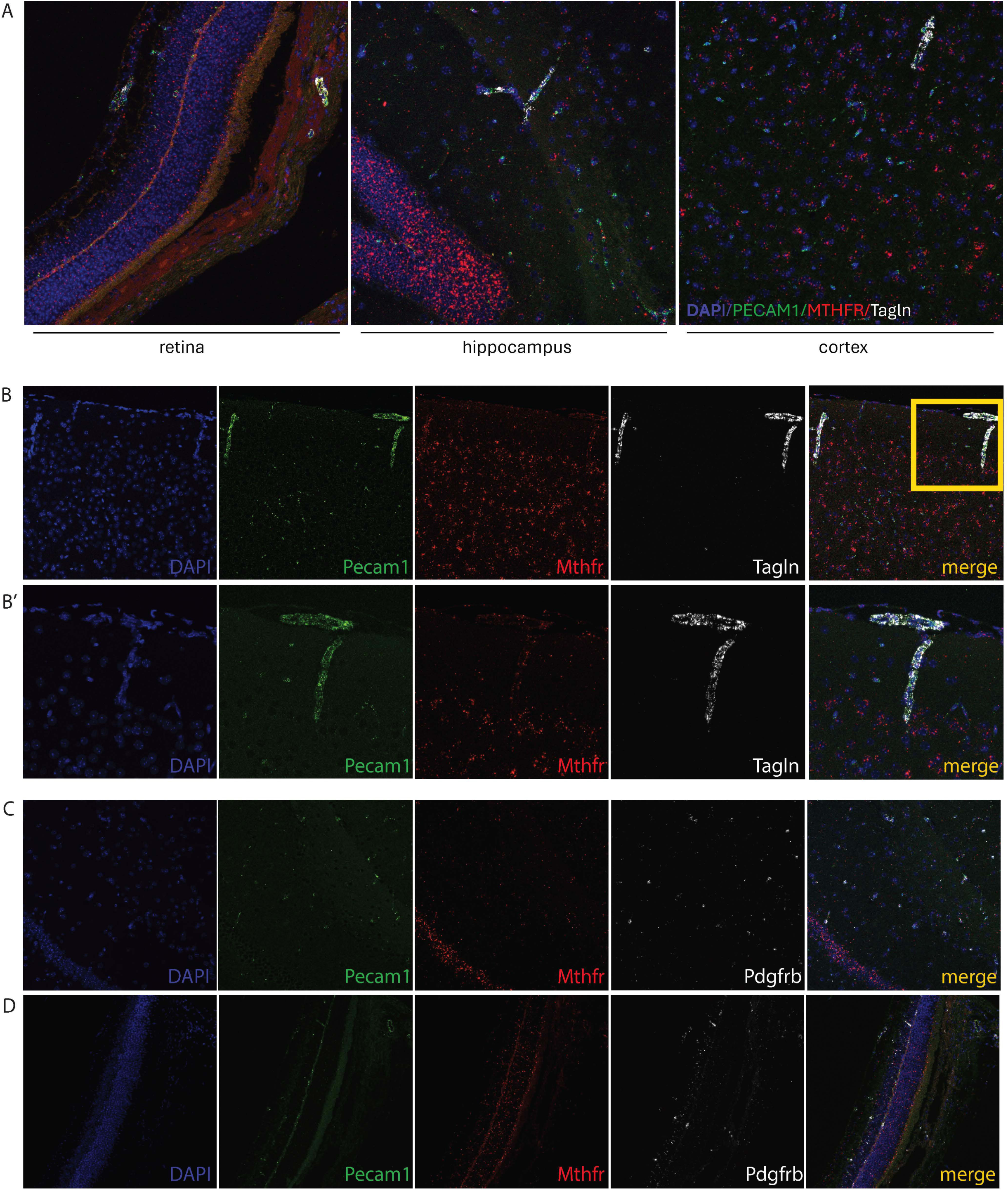
*Mthfr* is widely expressed in brain and retina. Representative confocal images showing Mthfr expression in retina, hippocampus and cortex using RNAscope. Merged images show expression of nuclear DAPI (blue) endothelial *Pecam1* (green) and vascular smooth muscle *Tagln* (white) show overlapping expression with *Mthfr*. Images are taken at 20x. (Figure 5A) Additional cortical vessels showing overlapping *Mthfr* and vascular gene expression at 20x (Figure 5B) and 40x (Figure 5B’). Cortical vessels (Figure 5C) and retinal vessels (Figure 5D) showing overlapping *Mthfr* and pericyte *Pdgfrb* (white) expression at 20x. (n=3/sex/genotype.)

### 3.6. Shared differentially expressed proteins between brain and retina are primarily associated with mitochondrial dysfunction and the ubiquitin-proteosome system

We performed tandem mass tag proteomics on brains and retinas from 6-month *Mthfr^677C>T^* mice. Using principal component analyses to visualize proteomic data, we reveal that both sex and *Mthfr^677C>T^* genotype similarly contribute to variation in brain (Figure 6A). While sex also contributes to variation in retina, genotype shows a more pronounced contribution (Figure 6B). In brain, we identified 904 differentially expressed proteins (DEPs) between CC vs TT, and 147 discrete DEPs in CC vs CT, with 151 shared DEPs (Supplementary Figure 1A). We performed STRING analysis on the CC to TT comparison which identified the largest cluster of DEPs was primarily associated with synaptic function (Supplementary Figure 1C). DEPs were significantly enriched for gene ontology (GO) terms relating to synaptic translation, metabolism, and protein-RNA complex association (Supplementary Figure 1E). In retina, we identified 540 DEPs between CC and TT, 2 DEPs between CC and CT, and 74 shared DEPs (Supplementary Figure 1B). For the CC to TT comparison, the largest cluster of DEPs was primarily associated with acetylation using STRING analysis (Supplementary Figure 1D) alongside GO term enrichment of telomere organization and maintenance, DNA metabolic processes and NAD/NADH metabolic processes using emap analysis (Supplementary Figure 1F). We then compared the DEPs in brain and retinal datasets, finding 206 shared DEPs (Figure 6C). STRING analysis suggested many DEPs were associated with diverse metabolic functions (Figure 6D) and there was enrichment of protein sets associated with telomere maintenance and cell metabolism, but with the appearance of a cardiac cell function module (Figure 6E). The largest group of DEPs did not share a common STRING function and were marked “unknown”. However, this category includes genes of known function including those of the respiratory complex I (*Ndufv1*, *Ndufa4, Ndufa12*, *Ndufs1*, *Ndufs3*, and *Ndufb8*) mutations in which result in mitochondrial dysfunction. Shared DEPs from the “unknown” module were also differentially expressed in a vascular enriched proteomics dataset that compared AD patients to controls (Figure 6F).

**Figure 6.**
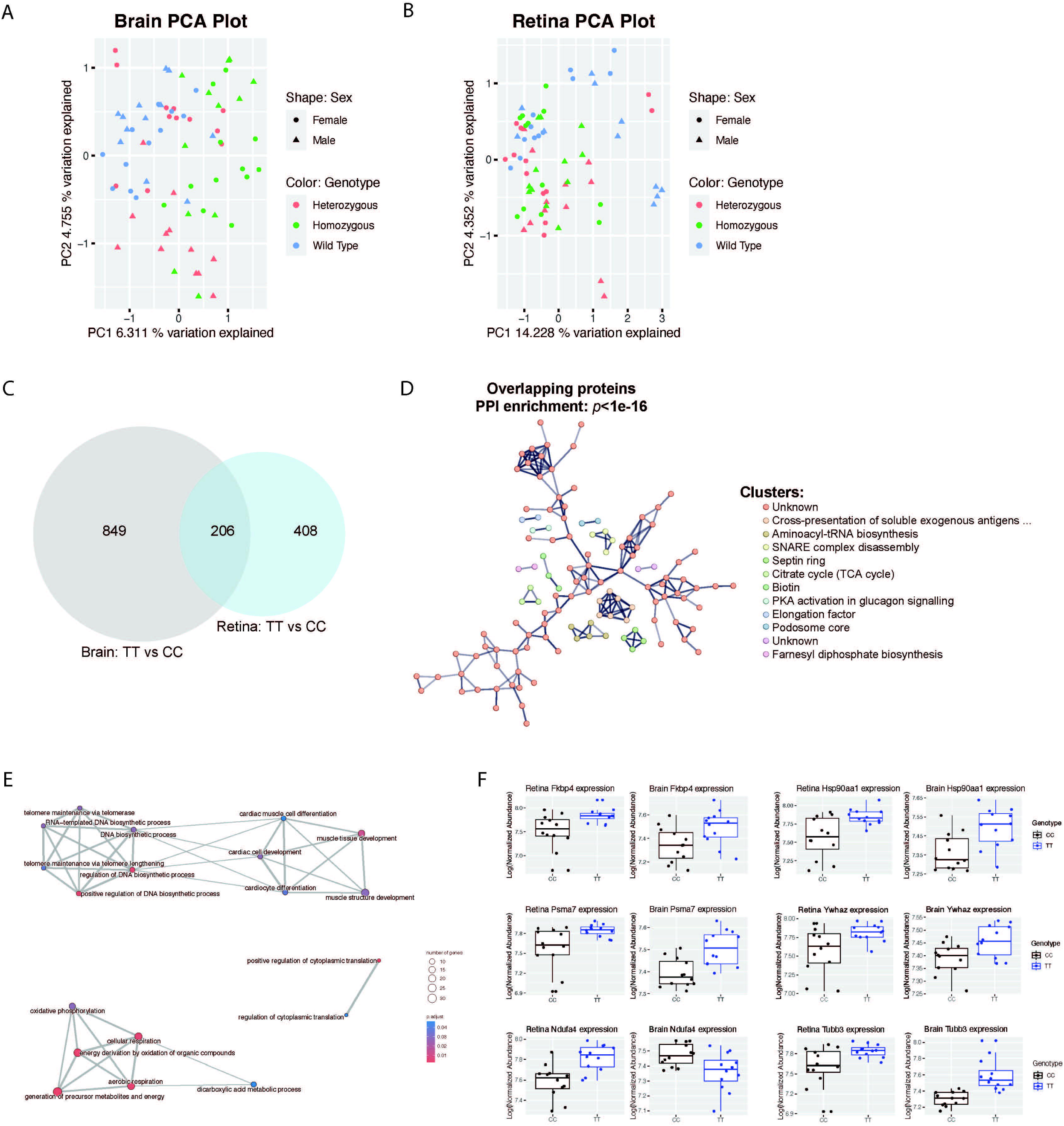
Differentially expressed proteins in brain and retina are primarily associated with mitochondrial dysfunction and the ubiquitin-proteosome system. Principal component analysis for brain (Figure 6A) and retina of *Mthfr^677C>T^* mice (Figure 6B). Venn diagram displaying overlap of DEPs from CC and TT brain and retina (Figure 6C). STRING analysis of the 206 DEPs summarizes the network of predicted associations for a particular group of proteins represented by nodes, and their predicted functional associations represented by the edges. (Figure 6D) Enrichment plot of “Biological Process” gene ontology terms (P < 0.05, FDR < 0.05) for the 206 DEPs, where p.adjust is the Benjamini–Hochberg adjusted P-value for the enriched ontology term and size of the node relates to the number of differentially expressed genes which belong to the enriched gene ontology term. (Figure 6E) Box and whisker plots of selected DEPs of interest. The average Log(normalized abundance) per mouse for each DEP is shown in CC and TT brains (left) and CC and TT retinas (right) for each DEP. (n=6 biological replicates/sex/genotype)

## 4. DISCUSSION

Phenotypes observed in *Mthfr^677C>T^* retinas mirror previous observations in *Mthfr^677C>T^* brains. Retinas demonstrate reduced vascular density and reduced vascular perfusion, with females showing more dramatic results. Additional vascular phenotypes suggest hypertensive conditions. Shared differential protein expression in retina and brain highlight pathways associated with the ubiquitin-proteosome system (UBS), cellular respiration, and cell survival/apoptosis, each of which may modulate vascular health. Because vascular contributions to ADRDs are becoming more widely accepted as hallmark pathologies and retinal vasculature may provide a direct physiological link to cerebrovascular health, our data supports the use of retinal vascular phenotyping as an ADRD biomarker.

The *MTHFR* gene codes for a key regulatory enzyme in folate and homocysteine metabolism. In physiological conditions it generates the folate derivative (5-methyltetrahydrofolate) that is necessary for homocysteine conversion to methionine. Methionine then generates S-adenosylmethionine (SAM), a critical donor of methyl groups for varied biological functions. Reduced MTHFR activity limits homocysteine conversion, which elevates homocysteine levels in the blood^63,64^. Elevated plasma homocysteine is associated with vascular damage in cardiovascular disease, with data demonstrating increased inflammation and endothelial cell dysfunction^65,66^. Cerebrovascular-relevant phenotypes have also been observed in mice on diets that induce hyperhomocysteinemia^67-69^.

The *MTHFR^677C>T^* polymorphism^70^ is associated with ADRDs as well as being independently linked to increased risk of peripheral vascular diseases including stroke^71^ and atherosclerosis^72^. Examining its role in neurodegenerative disease is broadly relevant because 20-40% of the global population is predicted to be either heterozygous or homozygous for *MTHFR^677C>T^*.^73^ *Mthfr* is expressed in all cells within the neurovascular unit, providing numerous opportunities for *Mthfr* to modulate signaling, but a molecular mechanism for vascular dysfunction remains unknown and until recently, no mouse model existed in which to study the polymorphism.

Retinal vascular pathologies often occur alongside microvasculature damage within other organs and may precede the progression of systemic vascular diseases. Retinal abnormalities including changes in vessel diameter and tortuosity, provide clinically accessible diagnostic elements for systemic diseases,^74^ including increased risk for diabetes,^75^ obesity,^76^ and cardiovascular disease^77^ as well as hypertension,^78^ stroke,^78^ and atherosclerosis.^79^ Each of these systemic diseases are independent risk factors for developing ADRDs, underscoring both the importance of vascular health in neurodegenerative disease and the usefulness of the retina as a biomarker. Here we show age-related reduction of retinal vascular density in female TT mice, and a reduced number of branching vessels in male and female TT mice, suggesting simplification of the vascular network. These phenotypes may contribute to a change in perfusion/oxygen availability for the surrounding tissue, priming it for damage once additional risk factors associated with aging and disease are introduced.

To further examine perfusion capability in the *Mthfr^677C>T^* retina, we measured the rate and intensity of fluorescein dye in the retinal vessels during *in vivo* angiography. Complementing the vessel density data, we observed reduced perfusion of fluorescein in TT mice, with older females showing the most notable reduction. In addition to changes in overall density, individual vessel morphology may also be contributing to reduced perfusion. *Ex vivo*, we noted TT retinas had narrowed arterioles and dilated venules, suggesting an intrinsic vascular response to altered blood pressure. Taken together, these data promote a hypoperfusion hypothesis, which could explain the loss of vessel density and network simplification. Hypoperfusion could be occurring through several mechanisms including: changes in upstream blood pressure, endothelial cell and/or mural cell thickening resulting in reduced luminal diameter, and reduced ability of endothelial and/or mural cells to respond to dilatory cues.

Our observation of arteriovenous crossing events and vessel tortuosity in TT retinas further suggests dysfunctional perfusion. Both phenotypes are affiliated with chronic hypertension, which itself is affiliated with narrowed arteries and hypoperfusion. AVC increases the odds for ischemic events, and vessel tortuosity can occur in response to an ischemic environment.^80-82^ These phenotypes are commonly seen in clinical imaging, further suggesting that observations made in the mouse retinal vasculature may provide human-relevant comparisons.

To understand how the *Mthfr^677C>T^* variant might be specifically affecting vascular health, and to determine if the mechanisms of vascular dysfunction are shared between brain and retina, we first determined which cell types are expressing *Mthfr* in both brain and retina using RNAscope. We demonstrate that *Mthfr* expression is widely distributed in both tissues of *Mthfr^677C>T^* mice and colocalizes with known vascular cell markers for endothelial cells, vascular smooth muscle cells and pericytes. However, because of its widespread expression, we cannot predict a cell type driving the observable phenotypes. MTHFR deficiency could influence vascular health either through vascular cell intrinsic mechanisms or through extrinsic mechanisms in which *Mthfr*-expressing glia or neurons are acting on vascular cells, altering their function. While these data provide more information about the expression of *Mthfr* in the CNS, it will be necessary to use a conditional knock-in model and additional proteomic strategies in future studies to examine cell-specific deficiency.

To further elucidate mechanisms driving vascular damage in the brain and retina of *Mthfr^677C>T^* mice, we performed bulk proteomics separately on brain and retina tissue comparing 6-month CT or TT mice to control CC mice. STRING and GO term enrichment analyses of DEPs between CC and TT brains suggest that MTHFR is regulating mitochondrial metabolism and synaptic translation. STRING and GO term enrichment analyses of DEPs between CC and TT retinas suggest that MTHFR is again regulating mitochondrial metabolism as well as telomere maintenance, suggesting an altered response to cellular aging. A total of 206 DEPs were shared between brain and retina, with the largest group of DEPs being assigned to an “unknown” gene ontology category. Further analysis of this group shows an association with mitochondrial dysfunction, including proteins that comprise Respiratory Complex I, the first large protein complex of the electron transport chain that is essential for the normal functioning of cell metabolism. Mutations in its subunits, including the DEPs Ndufv1, Ndufa4, Ndufa12, Ndufs1, Ndufs3, and Ndufb8, can result in a wide range of inherited neuromuscular and metabolic disorders. Defects in this complex are the most common in the oxidative phosphorylation disorders, and can result in mitochondrial dysfunction, oxidative stress, and are implicated in hypertrophic cardiomyopathy as well as increased damage from ischemia/reperfusion following an infarction.^83,84^ Additional shared DEPs categorized as “unknown” are associated with protein folding and trafficking. HSP90AA1 and FKBP4 have been associated with misfolded proteins in ADRDs^85^ and show increased expression in TT brains and retinas. Interestingly, a recent study using vascular enriched proteomics shows increased expression of HSP90AA1 and FKBP4 in Alzheimer’s patients compared to controls^86^ suggesting that TT vasculature may share human-relevant vascular cell profiles.

Several proteins in the 14-3-3 family were also differentially expressed, with 14-3-3ζ (also known as YWHAZ) particularly representing a key component in cell survival regulation. It interacts with many apoptotic proteins, including Raf kinases, BAX, BAD, NOXA and caspase-2, negatively regulating their activity. In this way it serves to protect the cell from environmental stresses, notably including hypoxia. 14-3-3ζ is highly conserved, and dysfunction has been implicated in neurodegenerative diseases, including Alzheimer’s disease.^87^ The enrichment analysis identified DEPs associated with cardiac/muscle cell gene ontology cluster. Because no cardiac cells were profiled, these likely represents proteins associated with regulation of vascular smooth muscle (VSM) and extracellular matrix (ECM) in the central nervous system. Altered smooth muscle cell function could contribute to chronic hypertension, resulting in the phenotypes we see in *Mthfr^677C>T^*retina and brain. Collectively, these data support our hypothesis that the same processes are driving vascular dysfunction in the brain and retina, and identify mechanisms for testing including mitochondrial dysfunction, hypoxia and impaired regulation of BBB components such as VSM and ECM.

Many factors, including genetics, contribute to age-related neurodegenerative disease. While some genetic risk is causative, other genetic and environmental factors may result in more subtle, additive damage over time. We hypothesize that vascular dysfunction occurring due to *Mthfr^677C>T^*may be sub-pathological but contributes to increased risk with age and onset of additional pathological features. Our study aimed to build on our existing work in the brain and characterize retinal vascular changes resulting from the 677C>T variant to both understand its function in the context of the retina and to determine the use of retinal vascular phenotypes to predict cerebrovascular health. Subsequent studies will include the addition of amyloid and tau pathology in the *Mthfr^677C>T^* model to induce neuronal stress alongside vascular stress.

## ACKNOWLEDGEMENTS

We would like to acknowledge the Mass Spectrometry and Protein Chemistry Service within Protein Sciences at The Jackson Laboratory for their contribution to the mass spectrometry work performed in this study. We gratefully acknowledge the contribution of Nick Gott and Ashley Dean and the Histopathological Service at The Jackson Laboratory for expert assistance with the work described in this publication.

We also thank Aleksandra Wojtas and Nicholas Seyfried for their help interpreting the human proteomics dataset from their recent publication.

This work was funded in part by an anonymous donation and the Diana Davis Spencer Foundation. Gareth Howell also currently holds the Diana Davis Spencer foundation chair for glaucoma research.

## AUTHOR CONTRIBUTIONS

A.M.R and G.R.H designed the study. A.M.R. performed animal harvests, tissue collection, and the associated experiments and data analysis. T.L.C. performed in vivo retinal imaging. M.M. analyzed proteomic data. A.M.R and G.R.H wrote the manuscript.

## CONSENT STATEMENT

No human subjects were used in this study.

## FIGURE LEGENDS

**Supplementary Figure 1. Mice carrying the *Mthfr^677C>T^* variant show differential protein expression in brain and retina.**

Venn diagram displaying overlap of DEPs from CC and TT brain (Supplementary Figure 1A) and retina (Supplementary Figure 1B). STRING analysis demonstrating pathways associated with protein-protein interactions for brain DEPs (Supplementary Figure 1C) and retina DEPs (Supplementary Figure 1D). Emap analysis demonstrating pathways associated with enriched GO terms for brain DEPs (Supplementary Figure 1E) and retina DEPs (Supplementary Figure 1F). (n=6 biological replicates/sex/genotype)

