## Supplementary figures and images for "Retinal vascular dysfunction in the *Mthfr^677C>T^* mouse model of cerebrovascular disease"

### Supplementary Figure 1

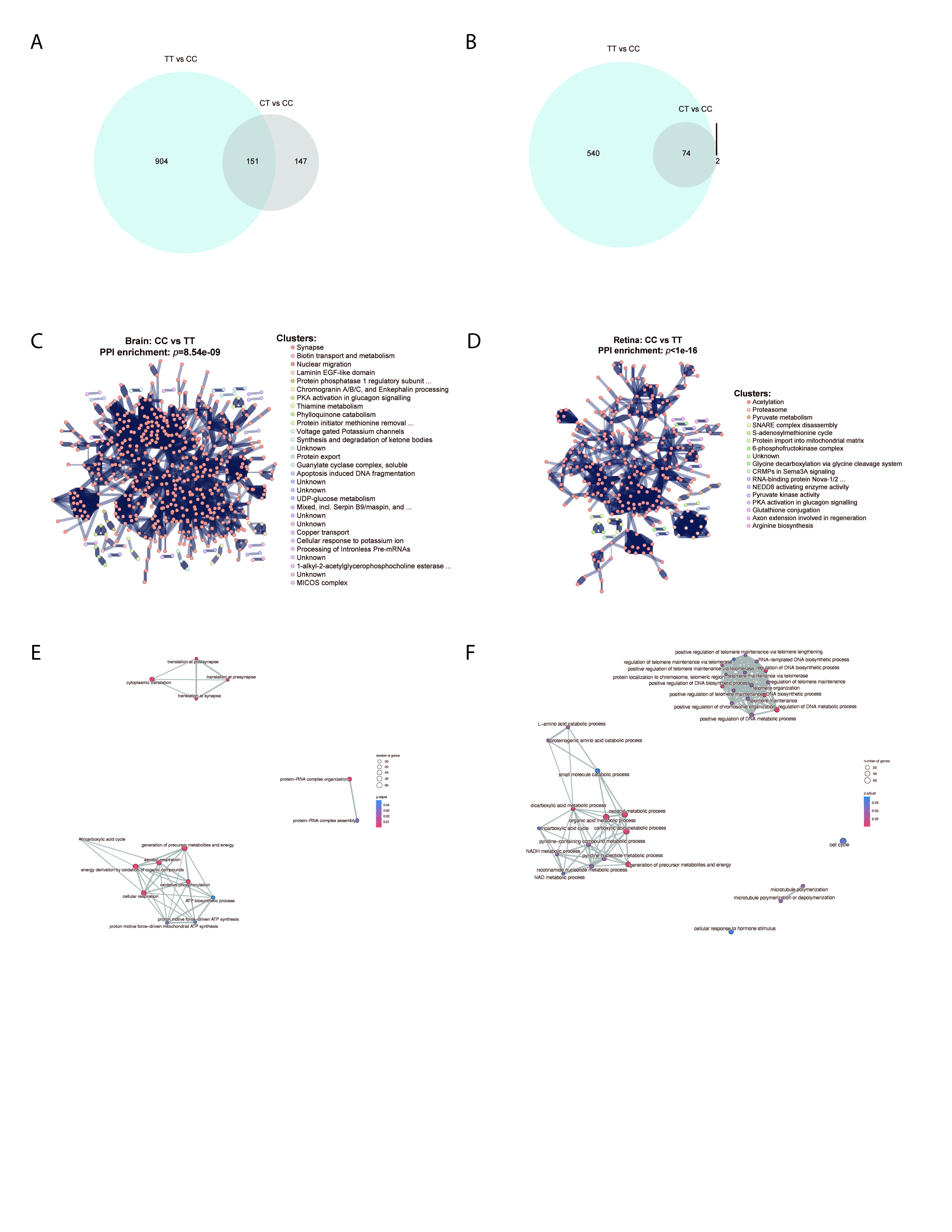
